# Plekha7 promotes retina organization and inhibits inflammation and photoreceptor loss

**DOI:** 10.64898/2026.04.03.716333

**Authors:** Lisa M. Cooper, Jiaxiang Chen, Ruifeng Lu, Ryan W. Feathers, Wan-Hsin Lin, Panos Z. Anastasiadis

## Abstract

Plekha7 associates with p120 catenin and localizes at apical cell-cell junctions where it has been implicated in junction stabilization. However, it’s in vivo function and underlying mechanisms of action remain unclear. Here, we show that constitutive Plekha7 loss in mice disrupts retinal organization, reduces retinal thickness, and induces RPE multinucleation, microglia infiltration, and photoreceptor loss. Plekha7 deficiency reduces the accumulation of cadherin–catenin complexes at apical junctions without altering overall cadherin levels, indicating a defect in junctional maintenance. Mechanistically, PLEKHA7 regulates cadherin trafficking by limiting levels of endocytosed E-cadherin, thereby maintaining junctional cadherin pools. Notably, these effects are largely independent of p120 catenin. In addition, further Plekha7 roles in the regulation of centrosome, cilia, microtubule dynamics, and cytokinesis, likely contribute to the phenotypic changes induced by Plekha7 knockout. These findings establish Plekha7 as a key regulator of cadherin trafficking and epithelial organization with relevance to retinal disease.

## Introduction

PLEKHA7 was initially described as a p120 catenin interacting partner and a component of cadherin-catenin complexes at the apical zonula adherens (ZA) of cultured epithelial cells. Early studies indicated a role in promoting junction maturation and suggested tethering minus ends of microtubules to the ZA through interaction with CAMSAP3 (Nezha) as a mechanism of action (Meng et al., 2008; Pulimeno et al., 2010). Recently, however, the recruitment of CAMSAP3 and microtubules to apical junctions was shown to be independent of PLEKHA7 and requiring the interaction of paracingulin (CGNL1) with ZO-1 at tight junctions (TJs) (Flinois et al., 2024). In turn, PLEKHA7 was shown to interact with PDZD11 to stabilize junctional nectins (Guerrera et al., 2016), to regulate calcium (Sluysmans et al., 2022) and copper homeostasis (Sluysmans et al., 2021) and to form a complex with tetraspanin 33 and ADAM10 to regulate susceptibility to staphylococcal α-toxin (Popov et al., 2015; Rouaud et al., 2020; Shah et al., 2018). Additionally, we reported that junctional PLEKHA7 suppresses the expression of proteins normally associated with proliferation, migration, and de-differentiation, via its association with the RNA interference machinery, including the RNA-induced silencing complex (RISC) (Kourtidis et al., 2017; Kourtidis et al., 2015). The PLEKHA7-recruited junctional RISC contains ∼30 miRNAs and 500 mRNAs and is responsible for the suppression of protein expression by PLEKHA7 (Kourtidis et al., 2017).

Despite its reported functions in junction stabilization and signaling, constitutive Plekha7^−/−^ knockout rats (Endres et al., 2014) or mice are viable, reproduce normally, and exhibit subtle phenotypes involving blood pressure regulation (Endres et al., 2014) and susceptibility to S. aureus α-toxin (Popov et al., 2015), respectively. The lack of overt epithelial pathologies suggested that Plekha7 is not an essential junctional component *in vivo* (Endres et al., 2014). Nonetheless, Plekha7 family members, including Plekha5 and 6, share the ability to interact with PDZD11, localize at cell-cell junctions, and may compensate for Plekha7 loss in the knockout mouse (Kourtidis et al., 2022; Sluysmans et al., 2021). Suggesting important roles in specific tissues, pathogenic variants of *PLEKHA7* have been associated with many human disorders, including hypertension (Levy et al., 2009), glaucoma (Khor et al., 2016; Vithana et al., 2012) and several eye syndromes, cleft lip with or without cleft palate (Cox et al., 2018), blepharocheilodontic syndrome, as well as metanephric and renal adenoma (Malacards) (Rappaport et al., 2013). Importantly, mutations in the E-cadherin/p120 complex, including pathogenic variants of *PLEKHA7* and *PLEKHA5*, cause mendelian non-syndromic cleft lip with or without cleft palate, indicating an important functional relationship between Plekha7/5 with the cadherin-p120 complex (Cox et al., 2018). In addition to a well-established relationship of *PLEKHA7* variants with primary angle closure glaucoma (Khor et al., 2016; Vithana et al., 2012), a recent GWAS study identified *PLEKHA7* as a genetic locus significantly associated (p= 6.6 × 10^−19^) with reduced macular thickness measured by spectral-domain optical coherence tomography using 68,423 participants from the UK Biobank cohort (Gao et al., 2019). The strong genetic associations of *PLEKHA7* pathogenic variants with eye disorders prompted us to examine in more detail the effects of Plekha7 knockout in the murine eye. Constitutive Plekha7 loss resulted in dysmorphic changes in the retina, reduced retina thickness, RPE multinucleation and infiltration by activated microglia, and photoreceptor loss. We uncover Plekha7 domains essential for membrane recruitment and increased ZA integrity, reveal a role for Plekha7 in E-cadherin trafficking, and also uncover novel roles for Plekha7 in cilia formation and/or maintenance, as well as cytokinesis, that likely contribute to the phenotypic changes induced by Plekha7 knockout.

## Results

### Plekha7 knockout promotes retina disorganization and reduces retina thickness

The generation of constitutive Plekha7^−/−^ (LacZ) knockout ICR/C57BL/6J chimeric mice targeting exon 8 of *Plekha7* (**Figure 1A**) was reported previously (Popov et al., 2015). qPCR analysis confirmed loss of *Plekha7* mRNA expression using probes spanning exons 12-13, or 15-16 of *Plekha7* (**Figure 1B**), while western blot analysis indicated loss of Plekha7 protein expression in the kidneys, lungs, and eyes of Plekha7^−/−^ mice (**Figure 1C**).

**Figure 1.**
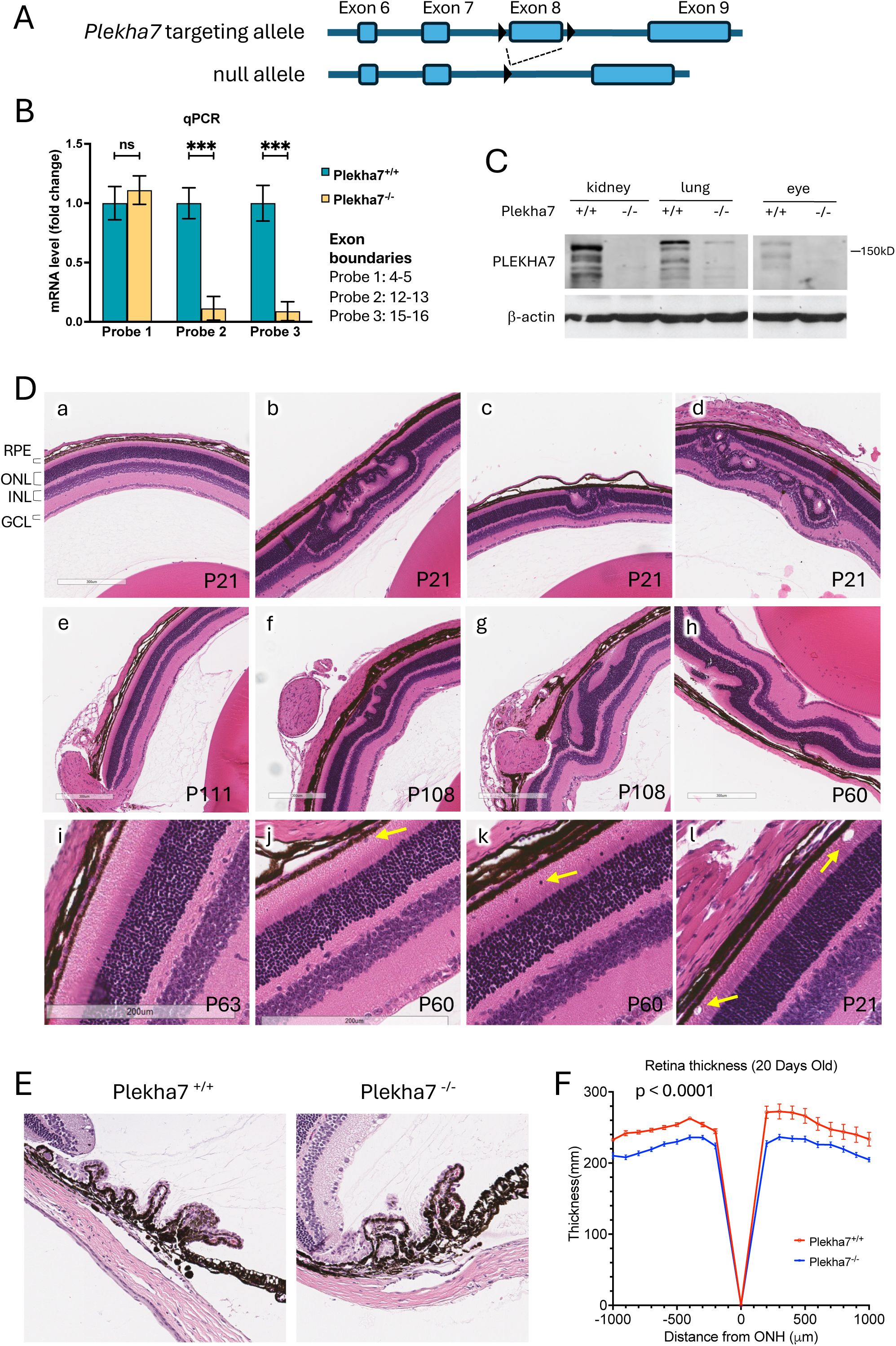
Plekha7 knockout promotes retina disorganization and reduces retina thickness. **A.** Schematic drawing depicting the generation of the Plekha7 constitutive knockout mouse. **B-C.** Confirmation of the Plekha7 knockout by RT-qPCR and western blot. In the *Plekha7^-/-^* mouse, Plekha7 mRNA expression is dramatically reduced (**B**) and the overall level of Plekha7 protein is lost in multiple tissues, including kidney, lung and retina (**C**). *** *p*< 0.001. **D.** Loss of Plekha7 causes retinal lamination defects at various timepoints. Paraffin embedded retinal sections stained with HCE. (**a**) wildtype mouse retina at p21 days, (**b-d**) Plekha7 knockout causes retinal lamination defects at p21 days, (**e**) wildtype mouse retina at p111 days, (**f-h**) Plekha7^−/−^ mouse retinal defects at p108 and p60 days, (**i**) wildtype mouse retina at p63 days, (**j-k**) nuclei of both pigmented and non-pigmented cells in the subretinal space of p60 Plekha7^−/−^ retinas (yellow arrow), (**l**) large inclusions in the RPE-photoreceptor interface of p21 Plekha7^−/−^ retina (yellow arrow). **E.** Paraffin embedded retinal sections stained with HCE. Plekha7 knockout does not induce gross morphological changes in the ciliary body. **F.** Plekha7 knockout reduces overall retina thickness at p20 (mean ± SEM, *n*=7, p < 0.0001). Thickness was measured at 0, 100, 200, 300, 400, 500, 600, 700, 800, 900, and 1000 μm from the optic nerve head (ONH) in the dorsal and ventral hemispheres, as indicated.

Plekha7^−/−^ chimeric mice were crossed with control C57BL/6J mice to generate heterozygous Plekha7^+/−^ mice, which were crossed together to generate Plekha7^−/−^ and Plekha7^+/+^ littermate controls. Postnatal retinas at different time points were then examined for potential defects in overall retina lamination and structure. In control retinas the four nuclear layers, retinal pigment epithelium nuclear layer (RPE), outer nuclear layer (ONL), inner nuclear layer (INL), and ganglion cell nuclear layer (GCL) were well laminated (**Figure 1Da, e, i**). However, Plekha7^−/−^ retinas exhibited abnormal retina organization. Locally disorganized regions were frequently detected in *Plekha7*-deficient retinas at P21, P60, and P108. Lamination defects such as retinal folds were frequent and prominent in *Plekha7*-deficient retinas and absent from retinas of control mice. These defects involved primarily the ONL, photoreceptor, and RPE layers (**Figure 1Db-d, f-h**).

Closer inspection of areas of normal lamination in Plekha7^−/−^ retinas revealed additional defects, including the presence of nuclei of both pigmented and non-pigmented cells in the subretinal space (**Figure 1Dj, k**), as well as disruptions adjacent to the RPE layer (**Figure 1Dl**), often associated with eosinophilic inclusions (Saksens et al., 2016). Gross anatomic changes were not observed by HCE staining in the ciliary body or the trabecular meshwork (**Figure 1E**). However, overall retina thickness was significantly smaller in Plekha7^−/−^ retinas compared to control at all time points tested (**Figure 1F, S1A**).

### Plekha7 is essential for ZA organization in the murine eye

In mature retina, Plekha7 localizes at the outer limiting membrane (OLM) and the RPE layers (Pulimeno et al., 2010). This localization is completely abolished in Plekha7^−/−^ retinas (**Figure 2A**). Similarly, in the ciliary body, a structure important for aqueous humor formation and intraocular pressure (Coca-Prados and Escribano, 2007; Pang et al., 2021a), Plekha7 localization at the apical junctions between pigmented and non-pigmented ciliary epithelial cells is completely lost in Plekha7^−/−^ cells (**Figure 2B**). As OLM is a major site of AJ and polarity protein accumulation and its disruption is associated with retinal lamination defects (Cho et al., 2012; Park et al., 2011), we then examined the effect of Plekha7 loss on the cadherin-catenin complex. The linear accumulation of N-cadherin, p120 catenin, and β-catenin at the OLM was disrupted and their overall expression, as assessed by IF, was reduced in both the OLM and the RPE cell-cell junctions of Plekha7^−/−^ retinas (**Figure 2C**, **S1B**). Knockout of Plekha7 exhibited an even stronger effect at the ciliary body, where the normal localization of N-cadherin and p120 at the apical ZA was completely abolished, while β-catenin was redistributed basolateraly primarily at the non-pigmented epithelial cells (**Figure 2D, S1B**). These results indicate an essential function for Plekha7 in ZA organization in the murine eye.

**Figure 2.**
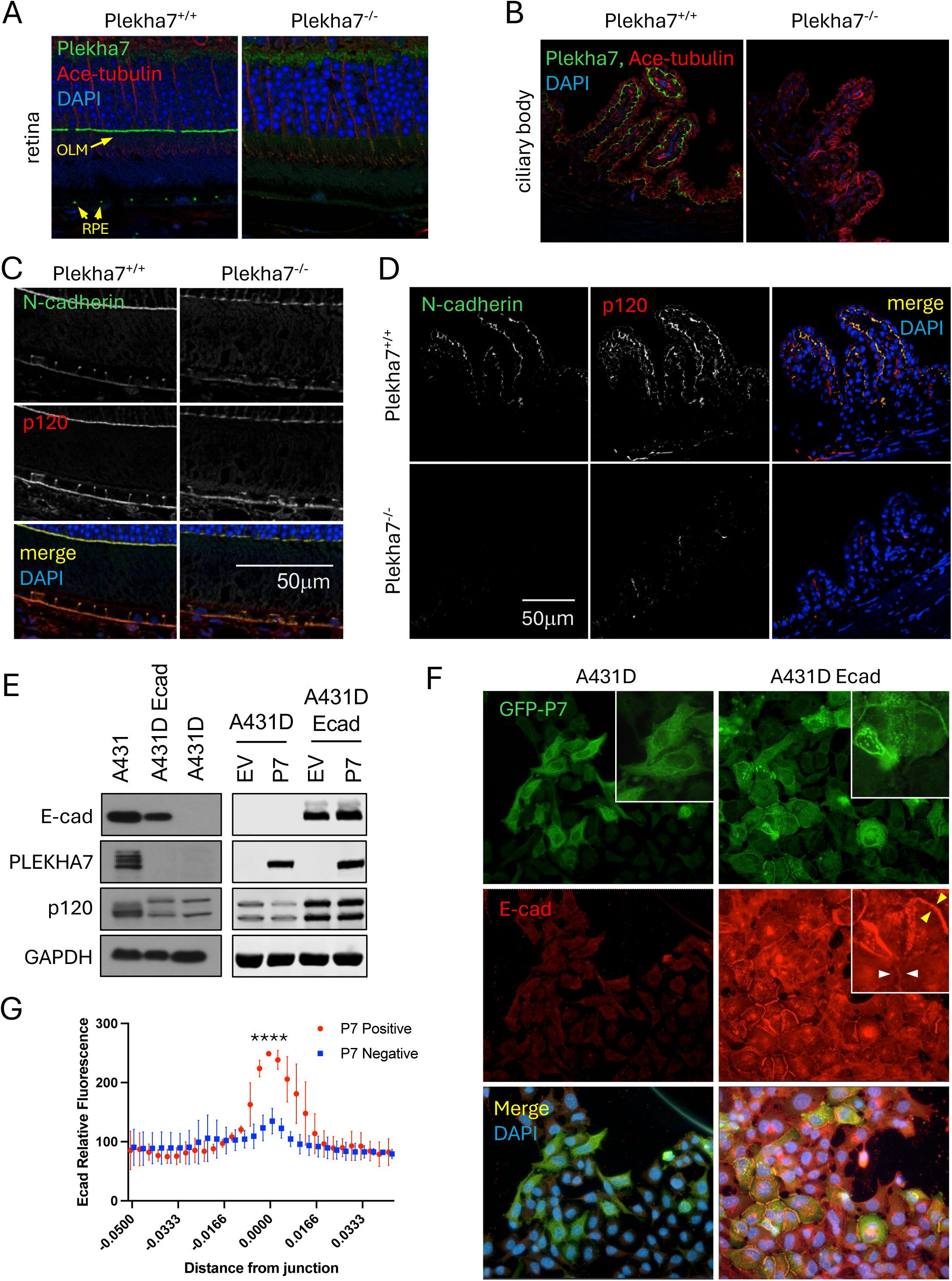
Plekha7 is essential for ZA organization in the murine eye. Immunofluorescence staining of paraffin embedded retinal sections. **A.** The predominant localization of Plekha7 (green) to the outer-limiting membrane (OLM) (yellow text) and retinal pigment epithelium (yellow text) is lost in Plekha7^−/−^ retinas. **B.** The exclusive localization of Plekha7 at the apical-apical interface of pigmented and non-pigmented epithelial cells in the ciliary body is lost in Plekha7^−/−^ retinas. **C.** Plekha7 knockout reduces the levels of N-cadherin and p120 catenin at the OLM and RPE. **D.** The localization of N-cadherin and p120 at the apical-apical interface of epithelial cell layers in the ciliary body is lost in Plekha7^−/−^ retinas. **E.** A431D cells lack endogenous PLEKHA7 expression (left panels) and ectopic expression of GFP-PLEKHA7 (P7) does not affect levels of exogenous E-cadherin (right panels). **F-G.** GFP-PLEKHA7 localizes at adherens junctions in A431D cells stably expressing E-cadherin and increases cadherin junctional accumulation (*n*=6, *p* < 0.0001).

Next, we examined potential vascular or other defects in Plekha7^−/−^ retinas. Co-staining of retina whole mounts with endothelial markers Caveolin-1 (Cav-1) and CD31 revealed no changes in the number of retinal arteries in knockout mice (**Figure S2A-B**). Similarly, co-staining retinas for Cav-1 and α-smooth muscle actin (αSMA) revealed no overt structural changes in the trabecular meshwork of Plekha7^−/−^ retinas (**Figure S2C**). Finally, co-staining lens epithelial cells with N-cadherin and p120 suggested that Plekha7 loss results in patterning defects, despite maintaining both N-cadherin and p120 expression and overall localization (**Figure S2D**).

### Plekha7 membrane recruitment is independent of p120 and regulates cadherin trafficking

To elucidate mechanisms of Plekha7 action we utilized A431D cells. A431D cells were derived from human A431 squamous carcinoma cells after prolonged dexamethasone treatment (Lewis et al., 1997) and lack all classical cadherins, express nectins (Troyanovsky et al., 2015), and express cytoplasmic p120 (Thoreson et al., 2000). Ectopic expression of E-cadherin recruits p120 to junctions and promotes cell adhesion, while expression of a p120-uncoupled E-cadherin mutant (764AAA; Δp120) fails to recruit p120 and promote adhesion strengthening (Thoreson et al., 2000). Unlike parental A431 cells, A431D cells lack expression of endogenous PLEKHA7 (**Figure 2E**). Similar to p120, recruitment of ectopically expressed GFP-tagged PLEKHA7 to A431D junctions depends on E-cadherin expression (**Figure 2F**). Furthermore, the recruitment of PLEKHA7 to junctions results in significant upregulation of E-cadherin accumulation at these sites (**Figure 2F, 2G**), consistent with Plekha7 effects in murine retina. Overall levels of E-cadherin are not affected by ectopic PLEKHA7 expression, suggesting that PLEKHA7 promotes the recruitment and/or stabilization of cadherin complexes at areas of cell-cell contact (**Figure 2E**).

Next, we ectopically expressed truncation mutants of PLEKHA7 to identify regions required for junctional localization and adhesion strengthening. A truncation mutant of PLEKHA7 containing the first 769 amino acids and lacking the C-terminal region (P7 1-769) was still able to localize properly at areas of cell-cell contact and induce junction strengthening (**Figure 3A-B**). In contrast, an internal deletion mutant lacking amino acids 538-769 (P7 Δ538-769) was selectively unable to localize at cell-cell junctions in A431D cells expressing E-cadherin and failed to increase junctional E-cadherin accumulation (**Figure 3A-B**). This region encompasses a proline rich domain (567-582) and the previously reported p120 interaction domain of PLEKHA7 (538-696)(Meng et al., 2008; Pulimeno et al., 2011).

**Figure 3.**
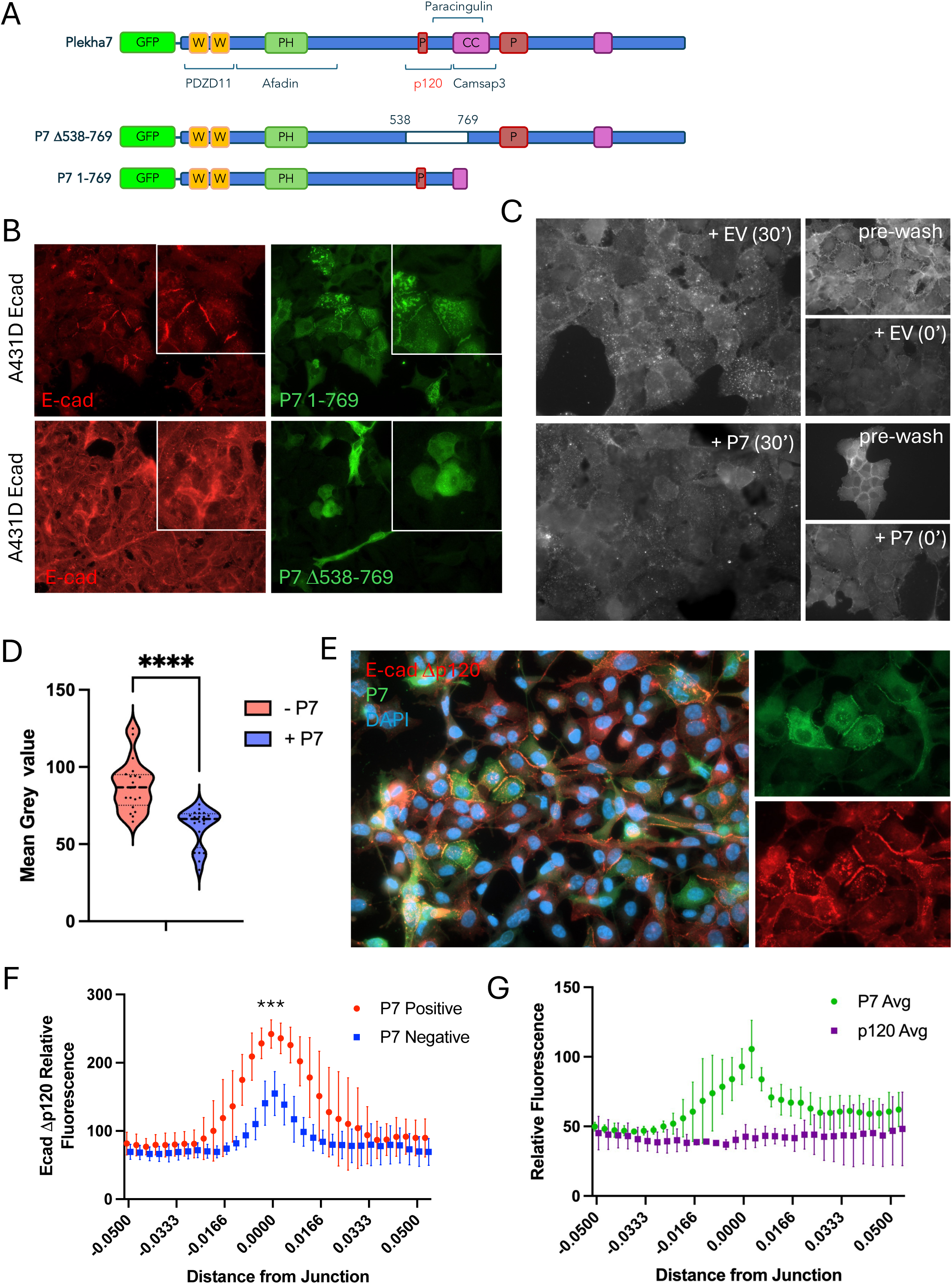
Plekha7 regulates levels of internalized E-cadherin and promotes E-cadherin junctional accumulation in a p120-independent manner. **A.** Schematic diagram of GFP-Plekha7 expression constructs indicating previously reported protein interaction domains. **B.** Unlike the GFP-P7 1-769 mutant, GFP-P7 Δ538-769 is unable to localize to cell-cell junctions and promote accumulation of junctional E-cadherin. **C-D.** Reduced accumulation of extracellularly labelled, internalized E-cadherin in A431D E-cad cells stably expressing PLEKHA7 compared to cells expressing empty vector (EV) after 30’ incubation at 37°C (*n*=20, *p* < 0.0001, unpaired t-test). **E-G.** Transiently expressed GFP-PLEKHA7 localizes at adherens junctions in A431D cells stably expressing a p120-uncoupled E-cadherin mutant (E-cad Δp120) and significantly increases cadherin junctional accumulation in the absence of junctional p120 (*n*=6, *p* < 0.0001).

A well-established role of p120 is the regulation of cadherin endocytosis and trafficking resulting in increased cadherin accumulation at cell-cell junctions and adhesion strengthening (Davis et al., 2003; Ireton et al., 2002; Xiao et al., 2003). Therefore, we reasoned that PLEKHA7 association with p120 promotes its junctional recruitment and adhesion strengthening by regulating cadherin trafficking in a p120-dependent manner. In agreement, the accumulation of extracellularly labelled E-cadherin into the cytosol of cells stably expressing PLEKHA7 (after 30’ incubation at 37°C) was significantly reduced compared to cells lacking PLEKHA7 expression (**Figure 3C-D**). To examine whether p120 is responsible for PLEKHA7 recruitment to A431D junctions we utilized A431D cells stably expressing a p120-uncoupled E-cadherin mutant (E-cad Δp120)(Thoreson et al., 2000). Exogenous PLEKHA7 was still recruited to areas of cell-cell contact (**Figure 3E**) and increased junctional E-cad Δp120 accumulation (**Figure 3F**) in these cells, despite the lack of p120 junctional localization (**Figure 3G**). Similarly, endogenous PLEKHA7 colocalized with E-cadherin at the junctions of SW48 cells, which lack expression of endogenous p120 (Ireton et al., 2002)(**Figure S2E**). Therefore, the data suggest a model whereby PLEKHA7 recruitment to apical adherens junctions and PLEKHA7-induced cadherin-catenin accumulation at sites of cell-cell contact are independent of p120 association.

### Plekha7 affects tubulin glutamylation and cilia assembly

Adjacent to the OLM, the inner segment (IS) contains the basal bodies, rootlets, and connecting photoreceptor cilia. Tubulin is highly glutamylated at the IS, a post-translational modification crucial for ciliogenesis (He et al., 2018; He et al., 2025) and overall structure of the photoreceptor cilium (Sergouniotis et al., 2014; Sun et al., 2016). Staining for glutamylated tubulin (Glu-tubulin) revealed a marked reduction of Glu-tubulin levels in the IS of Plekha7 knockout retinas compared to control, and significantly reduced thickness of the IS in retinas lacking Plekha7 expression (**Figure 4A-B**). Interestingly, significant overlap was observed in the IS of control retinas between Plekha7 staining and staining for centrin (Sergouniotis et al., 2014), a marker of centrosomes and basal bodies of cilia (**Figure 4C**).

**Figure 4.**
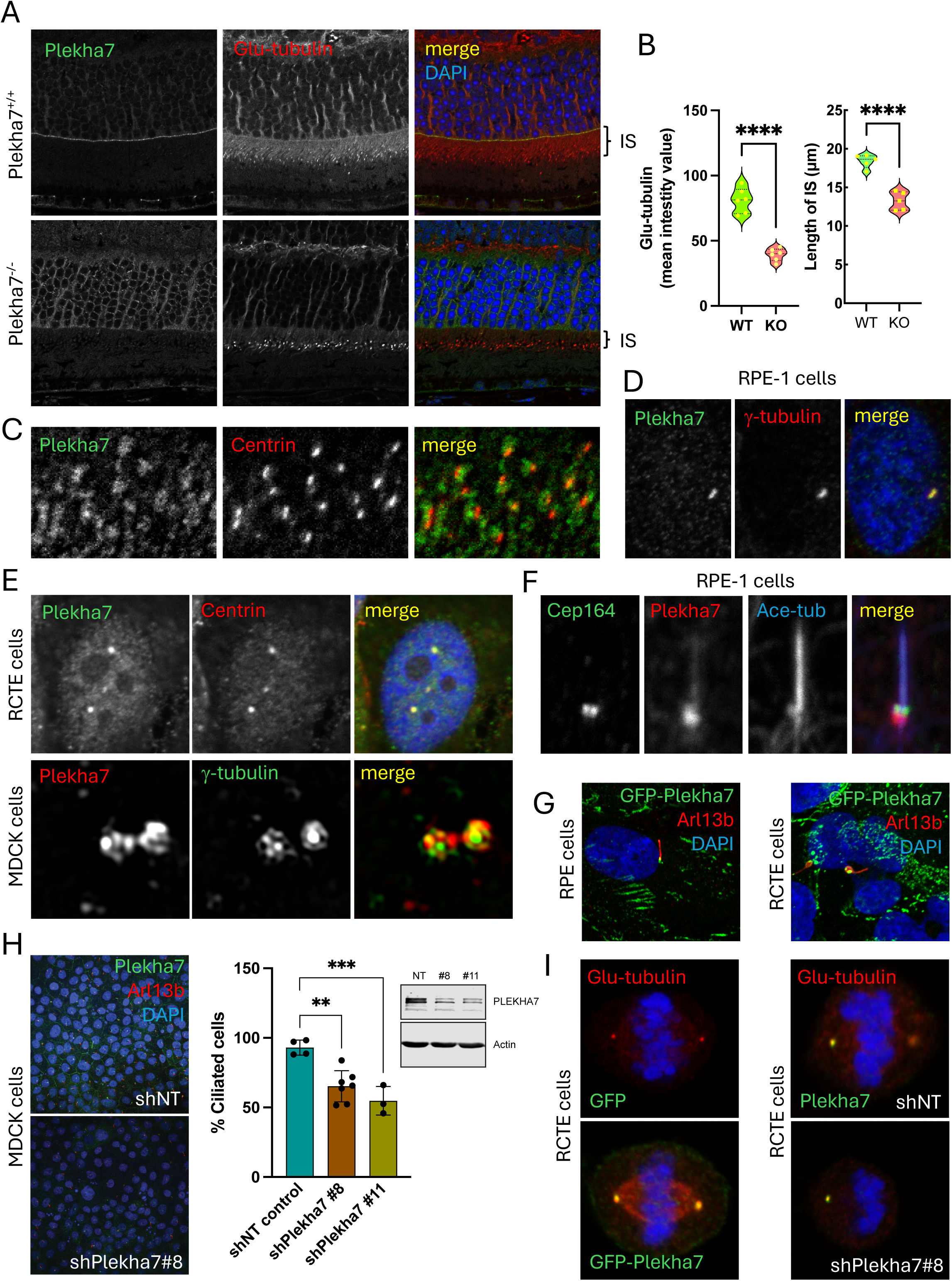
Plekha7 affects tubulin glutamylation and cilia assembly. **A-B.** Plekha7 knockout reduces overall glutamylated tubulin staining in the retinal inner segment (IS) and reduces IS length (*n*=6, *p* < 0.0001). **C.** Co-localization of Plekha7 with basal body marker Centrin in the mouse IS. **D.** Co-localization of Plekha7 with γ-tubulin at the centrosome of RPE-1 cells. **E.** Co-localization of Plekha7 with Centrin in RCTE cells (upper panels) and with γ-tubulin at the centrosomes of MDCK cells (lower panels, SR-SIM). **F.** Plekha7 localization at the basal body of quiescent RPE-1 primary cilia is distinct from the localization of Cep164, a specific marker of distal appendages. **G.** Ectopically expressed GFP-Plekha7 localizes to the basal body of primary cilia in RPE-1 and RCTE cells indicated by staining with the specific cilia marker Arl13b. **H.** shRNA-mediated knockdown of Plekha7 reduces the number of ciliated MDCK cells following serum deprivation (**p < 0.01, ***p < 0.001, one-way ANOVA with Dunnett’s multiple-comparison test). **I.** Ectopic expression of GFP-Plekha7 increases glutamylated tubulin (Glu-tubulin) in RCTE cells, while shRNA-mediated depletion decreases Glu-tubulin staining of mitotic spindles.

A centrosomal localization of PLEKHA7 was reported previously but was not further validated or explored functionally (Meng et al., 2008). Endogenous PLEKHA7 colocalized with γ-tubulin and/or centrin at centrosomes of immortalized human retinal epithelial (RPE-1) cells, normal human primary renal cortical epithelial (RCTE) cells, as well as canine epithelial MDCK cells (**Figure 4D-E, S3A**). Super-resolved structured illumination microscopy (SR-SIM) indicated the presence of PLEKHA7 in MDCK cell centrioles in close proximity to γ-tubulin (**Figure 4E**). Notably, the centrosomal localization of PLEKHA7 was also observed using GFP or Myc tag specific antibodies targeting ectopically expressed GFP- or Myc-tagged PLEKHA7 (**Figure S3B-C** and not shown). Examination of endogenous PLEKHA7 localization in RPE-1 cells indicated that it resides at centrosomes throughout the cell cycle (**Figure S3D**).

Interestingly, the junction-uncoupled Δ538-769 mutant of PLEKHA7 was still able to localize at centrosomes, indicating distinct mechanisms of recruitment to these sites (**Figure S4A**). As cells exit the cell cycle, mother and daughter centrioles move apically to become basal bodies of the apical cilium. Consistent with this, PLEKHA7 localizes at the basal bodies of cilia in serum-starved RPE-1, RCTE, and MDCK cells (**Figure 4F-H**). Depletion of endogenous PLEKHA7 in either MDCK or RPE-1 cells resulted in significant reduction of ciliated cells, indicating a role for PLEKHA7 in cilia formation and/or maintenance (**Figure 4H, S4B**). While the mechanism of this effect is beyond the scope of this study, tubulin glutamylation is essential for cilia extension (He et al., 2018; He et al., 2025). Interestingly, the level of Glu-tubulin during mitosis of RCTE cells was increased by ectopic expression of GFP-PLEKHA7 and decreased by PLEKHA7 depletion (**Figure 4I**).

### Junctional defects and multinucleation of RPE cells are induced by Plekha7 knockout

We utilized retina whole mounts to interrogate effects of Plekha7 loss in the RPE cell layer. In agreement with Figure 2A, the expression of Plekha7 was completely abolished in Plekha7^−/−^ RPE cells (**Figure 5A left**). Staining for p120 catenin revealed that the normal hexagonal pattern of RPE cell organization in control retinas was deregulated by Plekha7 loss, resulting in irregular RPE cell shapes, characterized by the presence of large multinucleated cells (**Figure 5A**). Apical actin staining was more diffuse and irregular in Plekha7^−/−^ RPE cells compared to the linear junctional staining in controls (**Figure 5A right**). Similar cell morphological defects were also observed by staining with the tight junction marker ZO-1 and were further quantified, indicating a significant increase in both large multinucleated RPE cells and small RPE cells in Plekha7^−/−^ retinas (**Figure 5B**).

**Figure 5.**
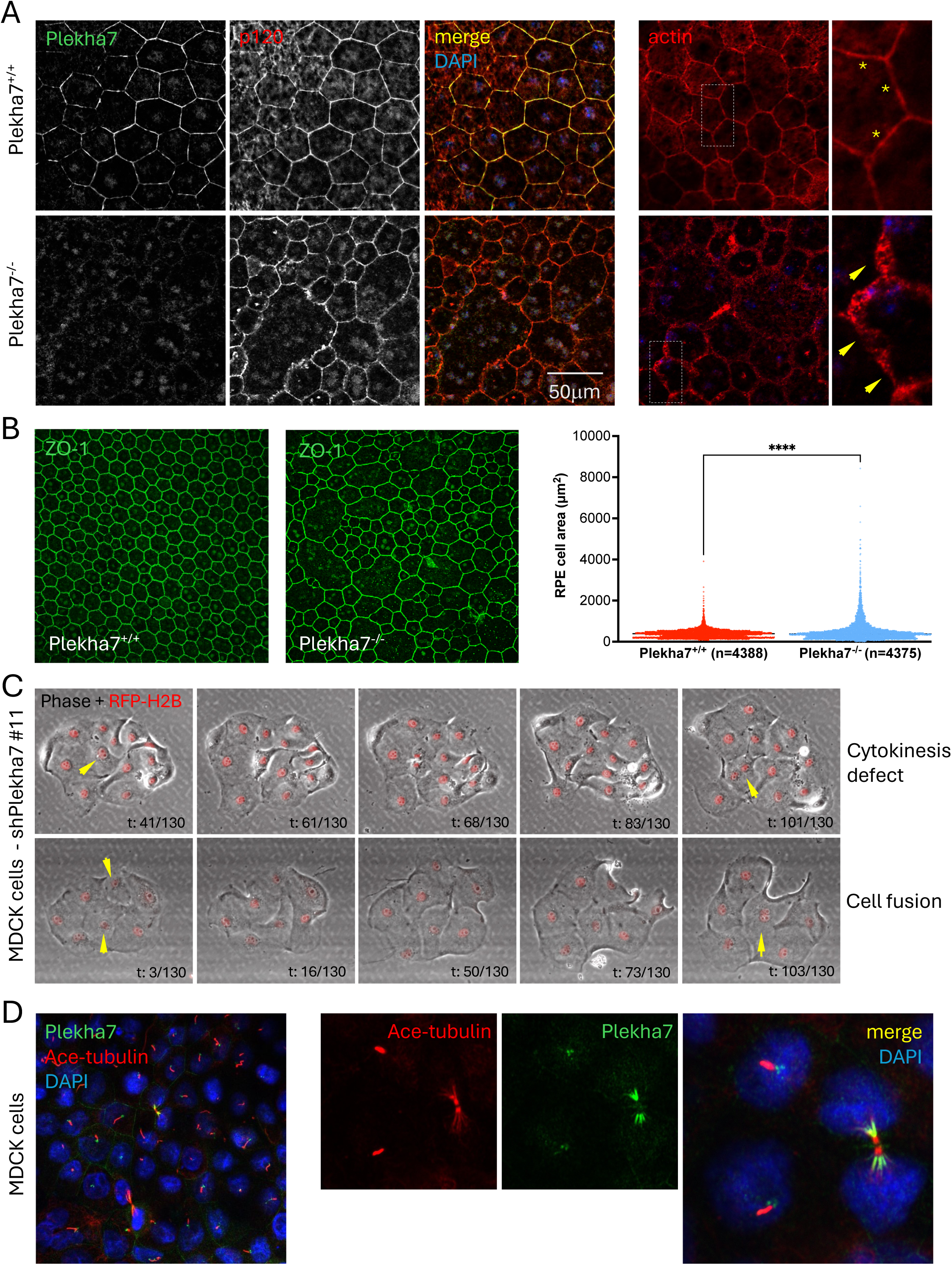
Junctional defects and multinucleation of RPE cells are induced by Plekha7 knockout. **A.** Knockout of Plekha7 disrupts the size and shape of retinal pigment epithelial cells (left panels), promotes multinucleation, and results in diffused actin localization at apical cell-cell junctions (right panels, arrows). **B.** Plekha7 knockout results in irregular size of retinal pigmented epithelial cells, as indicated by ZO-1 TJ staining (left panels) and cell area quantification (right, ****p < 0.0001). **C.** Representative images (n = 6 experiments) of Plekha7 knockdown MDCK cells expressing RFP-H2B and undergoing cell fusion, acquired by time-lapse microscopy (time in frames indicated below, 130 frames, 10 min interval). Yellow arrows indicate cell with cytokinesis defect (upper panel) or cells undergoing membrane fusion (lower panel). **D.** Co-localization of Plekha7 with acetylated tubulin at the midbody during anaphase.

Mechanistically, multinucleated cells may result from either cell fusion events forming large syncytia, or from cytokinesis defects (Patil et al., 2014; Saksens et al., 2016). We utilized MDCK cells stably expressing the nuclear marker RFP-H2B and live cell imaging to examine the potential effect of Plekha7 depletion on multinucleation. shRNA mediated depletion of Plekha7 resulted in multinucleation of MDCK cells that was common and happened either following cell division indicating a cytokinesis defect, or unrelated to cell division, indicating cell fusion (**Figure 5C**, **Movies 1, 2**). Interestingly, endogenous Plekha7 localized to the midbody during cytokinesis further suggesting a potential function in that process (**Figure 5D**).

### Plekha7 knockout promotes loss of photoreceptors

To detect possible defects in photoreceptor levels and organization we stained retinal sections with antibodies to cone Arrestin, a marker of cone photoreceptors, and Rhodopsin, which stains rod photoreceptor outer segments (Zhao et al., 2015). A marked reduction in the number of cone photoreceptors was observed in Plekha7^−/−^ retinas of age-matched mice (**Figure 6A**). In contrast, we observed no changes in the organization of Müller cells using vimentin as a specific marker (Hippert et al., 2021), or in the number and overall organization of bipolar rod cells stained for PKCalpha (PKCα; **Figure 6B**) (Ruether et al., 2010). Finally, we quantified overall cone photoreceptor numbers in three distinct regions of P360 retinal whole mounts: central, equatorial, and peripheral (**Figure 6C**). Progressively increasing photoreceptor loss was observed in equatorial and peripheral retinal regions.

**Figure 6.**
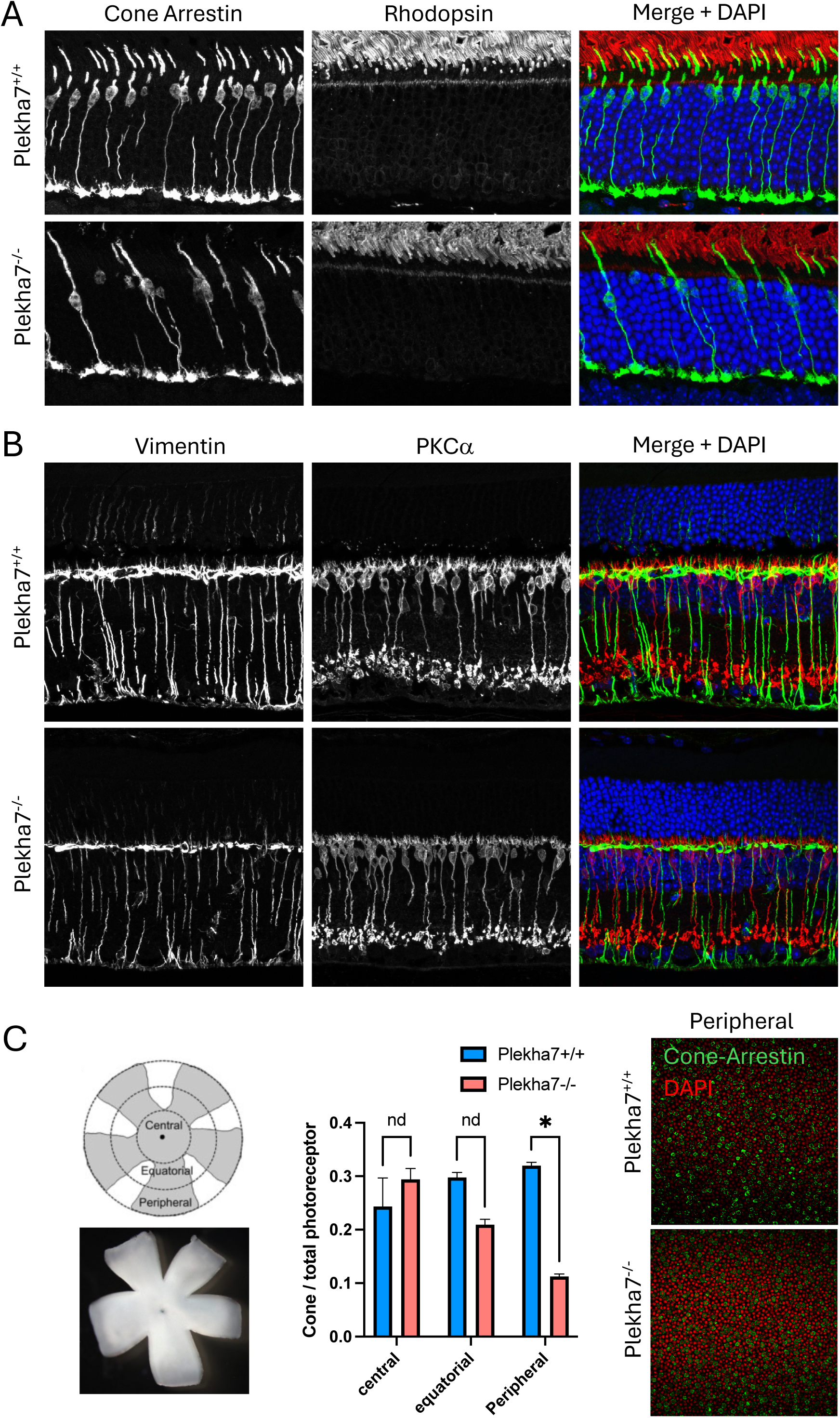
Plekha7 knockout promotes loss of photoreceptors. **A.** Plekha7 knockout results in reduction of cone photoreceptors at p360, as indicated by immunofluorescence staining for cone Arrestin. **B.** Knockout of Plekha7 does not affect the number of Müller cells, indicated by vimentin, or bipolar cells indicated by PKCα staining in the inner nuclear retinal layer. **C.** Schematic drawing of central, equatorial, and peripheral regions of the retinal whole mount (left). Quantification of cone photoreceptor distribution in different regions of Plekha7^+/+^ and Plekha7^−/−^ mouse retinas at p360 days shows significant reduction at peripheral regions (middle; mean ± SEM, *n*=4, \**p* < 0.05, two-way ANOVA with Dunnett’s multiple-comparison test). Reduced cone Arrestin staining at peripheral region of Plekha7^−/−^ mouse retinal whole mount (right).

### Loss of Plekha7 induces infiltration of activated microglia in the RPE layer

Several factors can promote photoreceptor loss, including IS defects and inflammation (Wright et al., 2010). Under physiological conditions, microglia reside in the OPL layer and are distributed in a mosaic-like pattern each exhibiting a small soma and extended ramified processes. However, under several pathological conditions, microglia undergo morphological changes characteristic of activation and infiltrate within the RPE and photoreceptor layers, where they contribute to photoreceptor degeneration (Combadiere et al., 2007; Ma et al., 2019; Ma et al., 2009).

The normal distribution and branched morphology of microglia in the OPL was also maintained in Plekha7^−/−^ retinas, as visualized by the selective microglial marker TMEM119 (**Figure 7A**). As expected, no infiltrating cells were detected in RPE whole mounts of control mice, stained for the general macrophage/microglia marker F4/80. In contrast, cell infiltration was evident in Plekha7^−/−^ RPE whole mounts (**Figure 7B**). Co-staining RPE whole mounts for TMEM119 and F4/80 indicated that infiltrating cells were indeed microglia (**Figure S4C**). Co-staining of agarose-embedded retinas with Iba-1 and F4/80, established microglia/macrophage markers, also indicated increased infiltration into the RPE and photoreceptor layers (**Figure 7C**, white arrows). Loss of dendrite arborization and increased soma size are characteristic morphological changes of activated microglia (Chen and Xu, 2015; Damani et al., 2011; Ma et al., 2019). In agreement, F4/80 staining revealed that RPE infiltrating cells exhibit morphology characteristic of microglial activation (**Figure 7D**). This was further quantified by determining the overall coverage of microglial dendrite projections, indicating a significant decrease in coverage between OPL resident and RPE infiltrating microglia (**Figure 7E**). Indeed, confocal z-sections confirmed that microglia intercalate within the Plekha7^−/−^ RPE cell layer and likely participate in active scavenging processes (Zabel et al., 2016) (**Figure S4D**, asterisks).

**Figure 7.**
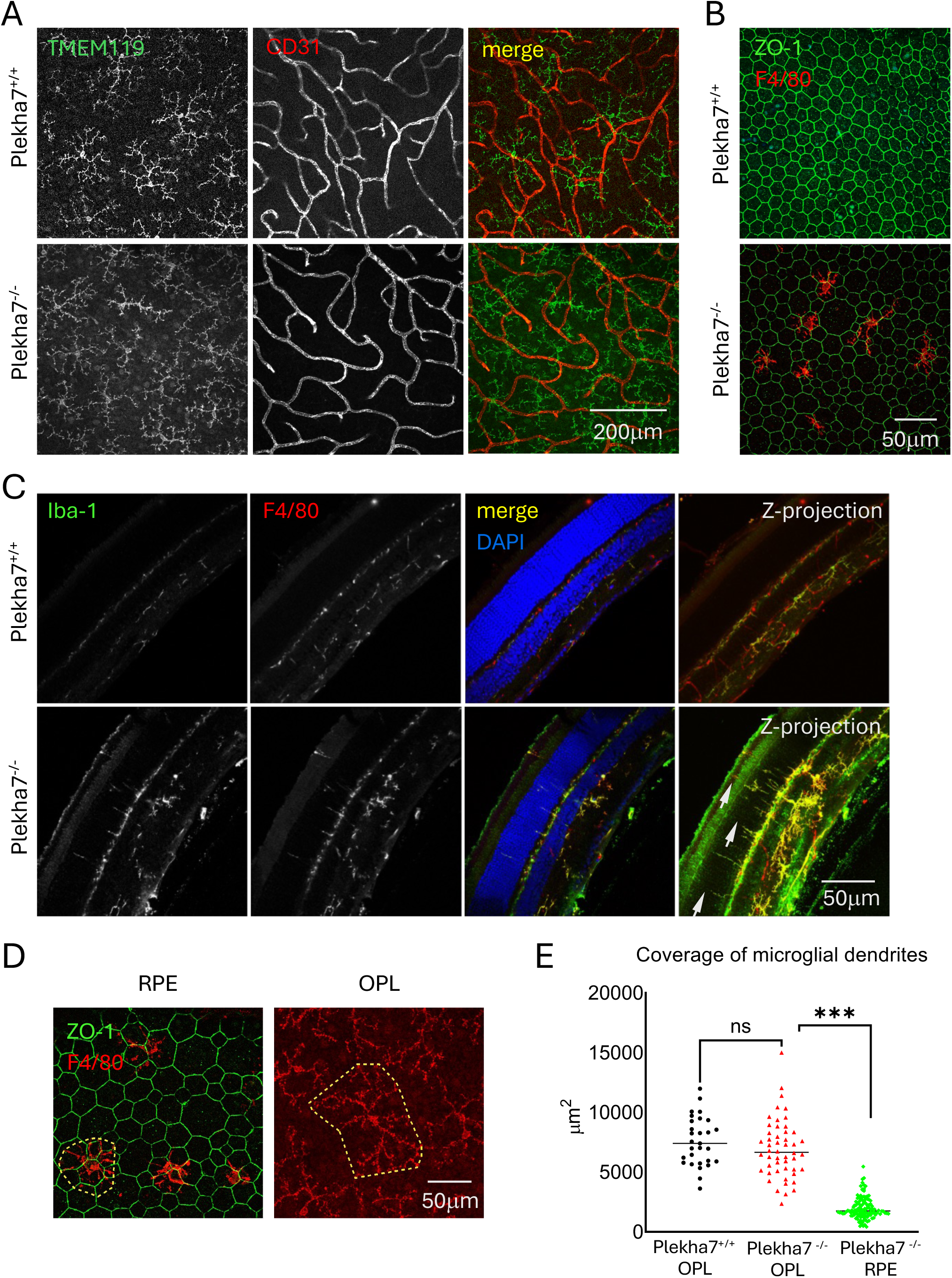
Loss of Plekha7 induces infiltration of activated microglia in the RPE layer. **A.** Microglia cells, indicated by TMEM119 staining (green), and vasculature, indicated by CD31 staining (red), exhibit normal distribution and morphology in the outer plexiform layer (OPL) of Plekha7^−/−^ retinas. **B.** Staining of RPE whole mounts with the general macrophage/microglia markers Iba-1 and F4/80 indicates cell infiltration in the Plekha7^−/−^ RPE layer. **C.** Co-staining of agarose-embedded retinas with Iba-1 and F4/80, established microglia/macrophage markers, indicate cell infiltration into the RPE and photoreceptor layers of Plekha7^−/−^ retinas (white arrows). **D.** F4/80 staining reveals reduced arborization and cell size of cells in the OPL and RPE of Plekha7^−/−^ retinas. **E.** Measurement of the total area covered by macrophage/microglia dendrites in the OPL and RPE layers indicates reduced size at the RPE of Plekha7^−/−^ retinas, while no infiltration was observed in the RPE of Plekha7^+/+^ retinas (ns, not significant, ***p< 0.001, two-way ANOVA with Dunnett’s multiple-comparison test).

## Discussion

Since its original introduction as an adhesion related protein, the specific function of PLEKHA7 at the junctions has been controversial. The main reason for this is that original studies in cultured epithelial cells supporting an important role for PLEKHA7 in promoting junction stability and maintenance (Kourtidis et al., 2015; Meng et al., 2008; Paschoud et al., 2014) have not been substantiated *in vivo*. Indeed, Plekha7 knockout rats and mice lack overt pathologies, while their phenotypes in attenuating salt-sensitive hypertension (Endres et al., 2014), or susceptibility to S. aureus α-toxin (Popov et al., 2015) were not directly correlated to Plekha7’s adhesive function but to regulation of intracellular calcium levels (Sluysmans et al., 2022), or to a complex formation with PDZD11, tetraspanin 33, and ADAM10, the α-toxin receptor (Rouaud et al., 2020). In this study, by using the murine eye as a model, we conclusively show that Plekha7 plays a crucial role in apical junction stability and/or maintenance.

In cultured epithelial cells, PLEKHA7 promotes accumulation of cadherin-catenin complexes at apical junctions and strengthens actin localization at the ZA (Kourtidis and Anastasiadis, 2016; Kourtidis et al., 2015; Meng et al., 2008; Paschoud et al., 2014). These effects were also evident in the murine retina, where Plekha7 loss significantly downregulated cadherin-catenin accumulation at apical junctions in the OLM and RPE layers and disrupted junctional actin organization. Surprisingly, the recruitment of PLEKHA7 to areas of cell-cell contact was independent of p120, although it required cadherin expression and a PLEKHA7 region previously reported to mediate p120 association. We postulate that an unknown protein recruits PLEKHA7 to cell junctions in a cadherin-dependent manner.

A recent study challenged the mechanism of PLEKHA7 action at the apical junctions which was thought to involve CAMSAP3 binding and MT minus end recruitment (Flinois et al., 2024). Instead, the recruitment of CAMSAP3 and MTs was shown to be independent of PLEKHA7 and required the interaction of paracingulin (CGNL1) with ZO-1 at tight junctions (TJs) (Flinois et al., 2024). We observed a significant reduction in endocytosed E-cadherin in cells expressing PLEKHA7 compared to control, which may suggest either decreased cadherin internalization from the cell membrane and/or increased trafficking back to the membrane. This effect likely accounts for the increased localization of cadherin complexes at apical junctions in the retina, as well as subsequent actin association and adhesion strengthening by Plekha7. Surprisingly, while E-cadherin endocytosis and junctional stability are thought to be regulated by p120, PLEKHA7-mediated cadherin accumulation at junctions appears to be largely independent of p120. Expression of exogenous PLEKHA7 increased the accumulation of p120-uncoupled E-cadherin at junctions, although to a lower extent than the accumulation of wild type E-cadherin (1.9-fold vs 3.4-fold over baseline, respectively; **Figures 2G, 3F**), suggesting that a higher order complex with p120 may further promote cadherin localization and function.

Interestingly, several retinal defects induced by Plekha7 loss were similar, albeit less severe, than defects induced by loss of apical polarity proteins including Crb1 (van de Pavert et al., 2004) and Pals1 (Cho et al., 2012; Park et al., 2011). For example, depletion of Pals1 in the murine retina promotes lamination defects, adhesion defects, photoreceptor loss, and microglia infiltration (Cho et al., 2012; Park et al., 2011). The ability of Pals1 to regulate cadherin trafficking (Wang et al., 2007) suggests a possible functional relationship between Plekha7 and the Crb/Pals1 complex that has not been explored to date.

A recent GWAS study identified *PLEKHA7* as a genetic locus significantly associated (p= 6.6 × 10^−19^) with reduced macular thickness (Gao et al., 2019). Consistent with this, knockout of Plekha7 reduced retina thickness across multiple age points with the most significant reduction observed in the IS layer. To visualize this layer, we stained for glutamylated tubulin, which was markedly downregulated in knockout retinas. Microtubule glutamylation is essential for cilia extension and the IS layer contains the basal bodies, rootlets, and connecting photoreceptor cilia (He et al., 2018; He et al., 2025; Sergouniotis et al., 2014; Sun et al., 2016). Furthermore, using immortalized RPE-1 cells, non-transformed MDCK cells, and primary normal RCTE cells, we validated the presence of endogenous Plekha7 in the centrosome and basal bodies of cilia, and detected a significant decrease in ciliated cells upon Plekha7 depletion. Interestingly, Plekha7 regulated the levels of glutamylated tubulin in the mitotic spindle, suggesting a function in the process of glutamylation and/or overall MT stability (Chen and Roll-Mecak, 2023). The data indicate a functional role for Plekha7 in centrosomes, cilia, and MTs that warrants further investigation.

A significant defect in RPE organization was observed in Plekha7 knockout retinas, with disruption of apical actin, irregular cell shape, and multinucleation. Plekha7 depletion in MDCK cells and live imaging indicated that both cytokinesis defects and cell fusion can account for this multinucleation phenotype. P120 catenin, a main Plekha7 interacting partner, has a known role in cytokinesis (van de Ven et al., 2016), and the localization of Plekha7 at the midbody argues for an important function for Plekha7. How Plekha7 may regulate cell fusion is unclear but may relate to regulation of membrane tension (Kim et al., 2015). Regardless of the mechanism of action, the observed defects in adherens and TJs argue for RPE barrier dysfunction, which may account for the increased infiltration of microglia. As Plekha7 can regulate the expression of a subset of mRNAs via a junctional RNA-induced silencing complex (Kourtidis et al., 2017; Kourtidis et al., 2015), it is also possible that depletion of Plekha7 results in expression of proteins that when secreted recruit microglia and promote their activation. Activated microglia are associated with retinal pathologies including photoreceptor degeneration (Fan et al., 2022; Ma et al., 2019; Zhao et al., 2015), as observed in the equatorial and peripheral regions of Plekha7 knockout retinas.

Overall, our findings establish Plekha7 as a critical regulator of apical junction integrity in the mammalian retina and uncover a previously unrecognized mechanism by which it controls cadherin trafficking to maintain junctional cadherin pools. Notably, this function occurs largely independently of p120 catenin, a major Plekha7 interacting partner and central regulator of cadherin stability. In addition to its role at the junctions, Plekha7 regulates centrosome and ciliary function, MT dynamics, and inflammation, linking junctional organization to broader aspects of epithelial architecture. Together, the results provide mechanistic insight into how Plekha7 dysfunction contributes to junction instability, inflammation, and tissue degeneration in the murine retina and suggest a potential role in human retinopathies. The presence of closely related family members may account for the relatively mild phenotypes observed in Plekha7 knockout mice (Kourtidis et al., 2022; Sluysmans et al., 2021), and further studies will be required to define their respective functions and contributions to human disease.

## Materials and methods

### Cell culture, constructs, antibodies

All cell lines used in this study were obtained from ATCC except RPE-1 (immortalized human retinal epithelial) and RCTE (normal human primary renal cortical epithelial) cells, which were provided by Dr. Jinghua Hu (Mayo Clinic, Rochester), and A431D cells, provided by Dr. Albert Reynolds (Vanderbilt University). Cells were used at low passage (<20) and tested negative for mycoplasma contamination. RPE-1 and RCTE cells were cultured in DMEM: F12 medium supplemented with 10% FBS, MDCKII cells in MEM medium supplemented with 10% heat inactivated FBS, HEK 293FT cells in DMEM medium supplemented with 10 % FBS, and A431D cells in DMEM medium supplemented with 10 % FBS and 1% penicillin/ streptomycin.

Full-length human PLEKHA7 (isoform 1, 1121aa) was cloned into the XbaI-HindIII sites of pcDNA3.1 Myc–6xHis vector. The GFP-PLEKHA7 construct, a gift from Dr. Masatoshi Takeichi (RIKEN, Japan (Popov et al., 2015)), was used as template to generate the GFP-P7 1-769, and GFP-P7 Δ538-769 constructs using specific probes and PCR amplification by Platinum™ SuperFi II Green PCR Master Mix (Themofisher). Amplified DNA fragments were digested with Xho1/BamH1 and cloned into the pEGFP-C1 vector (Clontech), sequence verified, and used in transient transfection experiments. GFP-PLEKHA7 constructs were also excised by NheI/BamH1 digestion and cloned into the pCDH-CMV-MCS-EF1α-Puro vector (System Biosciences, CD510B-1) before viral packaging and stable infection of A431D cells. GFP-Arl13b was a gift from Dr. Jinghua Hu (Mayo clinic, Rochester (He et al., 2018)) and pHIV-H2B mRFP was purchased from Addgene (#18982)(Welm et al., 2008). LZRS-wt-E-cadherin-neo, and LZRS-764-AAA-neo (E-cad Δp120) were described previously (Yanagisawa and Anastasiadis, 2006). All constructs were confirmed by sanger DNA sequencing. Lentiviral shRNAs were derived from the PLKO.1 based TRC1 (Sigma-Aldrich/RNAi Consortium) shRNA library. pLKO.1-puro Non-Target shRNA (SHC016) was used as control. Specific Plekha7 targeting sequences were shRNA Plekha7 #7: 5’-CGAATACCTGAAGTTGGAGAA-3’; shRNA plekha7 #8: 5’-CCTGAGTGCAAATAAAGAGAA-3’; and shRNA plekha7 #11: 5’-CTGAGCATCTTCTGTGAACAA-3’. Lentiviruses were produced in HEK293FT cells and used to infect cells as before (Kourtidis et al., 2017; Kourtidis et al., 2015; Lin et al., 2023). Infected cells were selected with puromycin for 48 hours and seeded immediately for subsequent analysis.

For detailed information on all the antibodies used in the study, see **Supplementary Table 1**. **Mice**

The constitutive Plekha7^−/−^ (LacZ) knockout ICR/C57BL/6J chimeric mouse was a gift from Dr. Masatoshi Takeichi (RIKEN Center for Developmental Biology, Kobe 650-0047, Japan) and obtained from Dr. Manuel R. Amieva (Department of Microbiology and Immunology, Stanford University School of Medicine, USA)(Popov et al., 2015). Plekha7^−/−^ chimeric mice were crossed with control C57BL/6J mice to generate heterozygous Plekha7^+/−^ mice, which were crossed together to generate age-matched Plekha7^−/−^ and Plekha7^+/+^ littermate controls. The following PCR primers were used for genotyping: WT forward: 5’-ACATGAACGCCTGGGTCAGG 3-’; WT reverse: 5’-GCCAGTGAGATGGTCCAGTT 3-’; KO forward: 5’-ATGGAAGGATTGGAGCTACG 3-’; KO reverse: 5’-GCCAGTGAGATGGTCCAGTT 3-’ (same as WT reverse). All animal experiments were approved by the Mayo Clinic Institutional Animal Care and Use Committee (Protocol number: A00004003-18-R24). Mice were housed in a pathogen-free environment with 12-hour light/dark cycle and *ad libitum* access to standard water and food.

Mice from different age-matched experimental and control groups were sacrificed and eyes were removed and fixed immediately. For paraffin sections, eyes were fixed first in Hartman’s Fixative (Sigma Aldrich, H0290) for 20 h, then in 4% paraformaldehyde in 1× PBS for another 20 h at room temperature, washed in 1× PBS four times (15 min each at room temperature), and dehydrated through a series of graded ethanol (30%, 50%, 70%, 85% and 95%; 1 h at room temperature each step), before paraffin-embedding. For agarose embedding fixed eyes were washed in 1× PBS four times (15 min each at room temperature) and then the optic nerve, cornea, iris, and lens were removed, and the remaining eyecup was submerged into a cryomold filled with preheated 6.5% low melting temperature agarose (68°C-70°C). The solidified agarose block was then sectioned using a vibratome into 100 μm thick slices that were mounted on cover slips and permeabilized overnight as reported (Pang et al., 2021b). For whole retinal pigment epithelial layer flat mount, eyes were fixed in 2% paraformaldehyde for 2 hours at room temperature. The anterior segment (cornea, iris, ciliary body and lens) was removed under an inverted microdissection microscope. Four to five radial cuts were made from the edge of the eyecup to the equator. Retinal tissue was removed and the remaining eyecups containing RPE, choroid and sclera were thoroughly washed and processed for immunofluorescence staining (Chen et al., 2016).

### Immunohistochemistry and immunofluorescence

For general morphological comparison, paraffin embedded eyes were cross sectioned (5 μm thickness) and stained with Hematoxylin and Eosin (HCE). Slide images were captured and analyzed using a ScanScope scanner and Aperio ImageScope software (Leica Biosystems). For retinal thickness measurement, HCE section images were taken at 20x magnification on either side of the optic nerve head using a Leica DM5000 microscope (Leica Microsystems, Wetzlar, Germany). ImageJ was used to measure retinal thickness at 100 μm intervals starting from the optic nerve head (ONH) and up to a distance of 1000 μm on each side. The whole retina thickness starting from the interface of the RPE layer and photoreceptor outer segment to the ganglion layer was measured. Three measurements were averaged at each point (Chen et al., 2020).

For immunofluorescence, paraffin embedded sections were dewaxed with xylene and subjected to antigen retrieval in citrate buffer (pH 6.0). Slides were treated with 3% H2O2 for 5 min to quench endogenous peroxidase activity and washed with 1x PBS solution containing 0.5% (w/v) Tween 20 (1x PBST), blocked in DAKO Serum-Free Protein Block (cat# X0909) and incubated with the indicated primary and secondary antibodies. Agarose embedded sections were permeabilized in 1x PBS containing 1% Triton-X 100 solution overnight, rinsed with 1x PBST (0.02% tween-20) three times, blocked with DAKO Serum-Free Protein Block (cat# X0909) and incubated with the indicated primary and secondary antibodies. RPE whole mount samples were permeabilized by methanol overnight at −20°C, rinsed with 1x PBST three times, blocked with DAKO Serum-Free Protein Block (cat# X0909) and incubated with the indicated primary and secondary antibodies.

For cell-based immunofluorescence, cells were seeded on glass coverslips and transfected (lipofectamine 2000, Invitrogen) 1 day later. 24-48 hours after transfection, cells were fixed in 1x PBS solution containing 4% paraformaldehyde and permeabilized in 1x PBS solution containing 0.2% Triton-X 100, then blocked in DAKO Serum-Free Protein Block (cat# X0909) and incubated with the indicated primary and secondary antibodies. Cell nuclei were visualized by DAPI (MilliporeSigma). Cells or tissues were mounted with ProLong™ Gold Antifade Mountant (Invitrogen™), immunofluorescence images were acquired on a LSM880 laser confocal microscope (ZEISS) and processed by ImageJ and Adobe Photoshop. In some cases, Z-series of images were acquired, and maximum intensity projection images were generated before data analysis. Junctional accumulation of proteins (E-cadherin, p120, PLEKHA7) was evaluated by perpendicular analysis of junctional intensity using ImageJ software (National Institutes of Health). Perpendicular lines were manually drawn across junctions (at least 10 measurements per field of view) and aligned based on the brightest point of E-cadherin. For super-resolution-structured illumination microscopy (SR-SIM) imaging, cells were cured for 60 h in a dark chamber before imaging. SR-SIM images were obtained using a Zeiss ELYRA S.1 SR-SIM with a 63× Plan-Apochromat 1.4 NA objective. SIM processing and channel alignment were rendered using ZEN imaging software.

### Cadherin Internalization

A431D cultures were seeded at 300K cells/well in a 12-well plate 24 h before experiments. At the start of the assay, cell medium was changed to Hanks’ balanced salt solution supplemented with 5 mM Ca^2+^ and 50 mg/ml bovine serum albumin and cells were allowed to equilibrate for 1 h at 37°C. Cells were then incubated for 1 h at 4°C with 1μg/ml HECD-1 IgG diluted in Hanks’ balanced salt solution. Coverslips were washed with ice-cold PBS to remove the unbound antibody, and cells were either fixed to visualize surface-bound antibody (pre-wash), subjected to acid wash (0.5 m acetic acid, 0.5 m NaCl; 3× 30 second washes) to remove surface-bound antibody (time 0’), or incubated at 37°C for 30 minutes in Hanks’ balanced salt solution before acid washing (time 30’). Cells were washed with PBS before fixation with 4% paraformaldehyde/ 0.12M sucrose for 20 min at 4°C, permeabilized with 0.2% TX-100 in PBS and incubated with secondary antibody for visualization. To assess protein internalization, the fluorescence intensity (mean gray value) of E-cadherin was measured in individual cells within the monolayer (with at least 10 cells analyzed per field of view) using ImageJ.

### Immunoblotting

Mouse tissues (whole eye, kidney and lung) were lysed in 1x RIPA buffer (50 mM Tris-HCl, pH 7.4, 150 mM NaCl, 0.25 % deoxycholic acid, 1 % NP-40, 1 mM EDTA, 1% SDS) with 1x proteinase (Protease Inhibitor Cocktail Set I, Roche) and HALT phosphatase inhibitors (Pierce). Mouse tissue lysate supernatants were collected following centrifugation at 20000-rpm for 20 min at 4 °C, and protein concentration was quantified (Pierce™ BCA Protein Assay Kit). Equal amounts of protein were subjected to SDS-PAGE and western blotting with appropriate primary and secondary antibodies (IRDye680 for mouse and IRDye800 for Rabbit). Western bolt images were acquired using an Odyssey CLx Infrared Imaging System (LI-COR biotech) and processed with Adobe Photoshop or Adobe illustrator.

Cultured cells were lysed in 1x Triton-X lysis buffer (150 mM NaCl, 1 mM EDTA, 50 mM Tris pH 7.4, 1% Triton X-100) containing 1x proteinase inhibitor (Protease Inhibitor Cocktail Set I, Roche) and HALT phosphatase inhibitors (Pierce), and lysates were handled as above for protein quantification and western blotting.

### qRT-PCR

Total RNA (2 eyes per animal) was isolated using TRIzol (Invitrogen) according to manufacturer’s protocol and further purified with the Qiagen RNeasy Plus Mini Kit (Qiagen 74134). RNA conversion to cDNA was performed using the high-capacity cDNA reverse transcriptase kit (Applied Biosystems). qPCR reactions were carried out using the TaqMan FAST Universal PCR master mix (Applied Biosystems) and the QuantStudio™ 6 Flex Real-Time PCR System (Applied Biosystems). Gene expression was quantified using the ΔΔCt method using normalization to GAPDH. The TaqMan assay IDs (Applied Biosystems, 4331182) used in this study: *GAPDH: MmSSSSSS15_g1*, plekha7, probe #1 (exon boundary 4-5): *Mm00555062_m1,* probe #2 (exon boundary 12-13): *Mm00555064_m1*, and probe #3 (exon boundary 15-16): *Mm01162828_m1*.

### Live-cell imaging

For live-cell imaging, images were captured by using an Olympus IX83 imaging system equipped with a Stage Top Incubator (Tokai Hit). To examine the mechanism of multinucleation upon Plekha7 knockdown, MDCK II cells expressing RFP-H2B and NT control or *Plekha7* targeting shRNA#11 were plated the day before imaging. Images were acquired every 10 min over a 20-24 hour period with a 20× phase objective. Phase-contrast and RFP fluorescence images were combined and used to identify cells undergoing multinucleation due to either cytokinesis failure or fusion. Movies were generated using ImageJ.

### Cilia

Cilia formation was induced by serum starvation as described (He et al., 2018; He et al., 2025). Briefly, RPE-1, MDCK II, or RCTE cells stably expressing GFP-Plekha7 were seeded on coverslips for 24 hours and then were serum starved (DMEM: F12 medium supplemented with 0.5 % FBS) for another 24 hours before fixation in −20°C methanol. For Plekha7 knockdown experiments, puromycin selected MDCK II cells expressing control or Plekha7 targeting shRNAs were seeded on coverslips, serum starved and fixed as above for immunofluorescence analysis.

### Statistical analyses

Statistical analysis was performed using the GraphPad Prism software. Statistical significance was determined by unpaired Student’s t test or by one-way or two-way ANOVA with Dunnett’s multiple-comparison test. Bar graphs present the data (mean ± SD or mean ± SEM) from multiple experiments or data points. Experimental details are indicated in the figure legends. P < 0.05 was considered statistically significant.

## ACKNOWLEDGMENTS

We are grateful for the time we spent with our late friend, colleague, and co-author of this study, Dr. Ruifeng Lu. He was a dedicated father and scientist and will be missed. We thank Brandy Edenfield and Dr. Laura Lewis-Tuffin for their technical support with immunohistochemistry and imaging, respectively. We also thank Dr. Jinghua Hu (Mayo Clinic, Rochester) for providing reagents and helpful discussions and Drs. Alan Marmorstein and Michael Fautsch (Mayo Clinic, Rochester) for helpful suggestions. The work was supported by Florida Department of Health James and Esther King Biomedical Research grant 24K01, and a Center for Biomedical Discovery (Mayo Clinic) grant (to PZA).

