## Supplemental figures and table for "Plekha7 promotes retina organization and inhibits inflammation and photoreceptor loss"

#### Supplementary Figures

**Supplementary Figure 1. A.** Plekha7 knockout reduces overall retina thickness at p136 (left; mean ± SEM, $n$=3, $p < 0.05$) and p60 days (right; mean ± SEM, $n$=5, $p < 0.0001$). **B.** Plekha7 knockout reduces the levels of β-catenin at the retina OLM and RPE (left panels). β-catenin localization in the ciliary body of Plekha7$^{-/-}$ retinas is diffused and excluded from the apical-apical interface of pigmented and non-pigmented epithelial cell layers (right panels).

**Supplementary Figure 2. A-B.** Plekha7 knockout does not affect vascular structure in the retina (ns, not significant) **C.** Immunofluorescence staining of endothelial cells (indicated by Cav-1) and myofibroblasts (indicated by αSMA) indicates no gross morphological changes in the structure of the trabecular meshwork at the ciliary body of Plekha7$^{-/-}$ retinas. **D.** Plekha7 knockout induces disorganization of lens epithelium (cell junctions indicated by N-cadherin and p120 staining). **E.** Co-localization of endogenous PLEKHA7 with E-cadherin at the cell-cell junctions of SW48 cells lacking p120 expression.

**Supplementary Figure 3. A.** Endogenous Plekha7 co-localizes with γ-tubulin at the centrosome of MDCK II cells. **B.** Ectopically expressed GFP-Plekha7 co-localizes with Centrin at the centrosome of MDCK II cells (GFP, green; Centrin, red). **C.** Ectopically expressed GFP-Plekha7 co-localizes with γ-tubulin at the centrosome of RCTE cells (GFP, green; γ-tubulin, red). **D.** Endogenous Plekha7 co-localizes with γ-tubulin at the centrosome of RPE-1 cells throughout the cell cycle.

**Supplementary Figure 4. A.** The junction-uncoupled PLEKHA7 Δ538-769 mutant localizes to the centrosome in RCTE cells. **B.** shRNA-mediated knockdown of Plekha7 reduces the number of ciliated RPE-1 cells following serum deprivation (****$p < 0.0001$, unpaired t test). **C.** Co-localization of TMEM119 and F4/80 cell markers in the RPE of Plekha7$^{-/-}$ retina whole mounts. **D**. Confocal z-sections of Plekha7$^{-/-}$ retina whole mount indicates F4/80 positive macrophage/microglia infiltrating the RPE cell layer (junctional ZO-1 staining). Cellular regions characteristic of active scavenging are highlighted with asterisks.

**<u>Supplemental Table 1</u>: List of antibodies used in this study.**

| Name | Source | Identifier |
|---|---|---|
| Rabbit anti-plekha7 | GeneTex | Cat # GTX131146 |
| Rabbit anti-beta actin | Cell Signaling | Cat # 4967L |
| Mouse anti-Acetylated tubulin | Sigma-Aldrich | Cat # T6793, clone 6-11B-1 |
| Rabbit anti N-Cadherin | Abcam | Cat # ab12221 |
| Mouse anti p120 ctn | Santa Cruz Biotechnology | Cat # sc-23873, clone 6H11 |
| Mouse anti Caveolin-1 | Santa Cruz Biotechnology | Cat # sc-894 |
| Mouse anti E-cadherin (HECD1) | Zymed/Invitrogen | Cat # 13-1700 |
| Goat anti PECAM-1 (M-20) (CD31) | Santa Cruz Biotechnology | Cat # sc-1506 |
| Mouse anti Polyglutamylated Tubulin (GT335) | AdipoGen | Cat# AG-20B-0020-C100 |
| Mouse anti Centrin | Millipore | Cat # 04-1624, clone 20H5 |
| Mouse anti Gamma Tubulin | Abcam | Cat# ab11316, clone GTU-88 |
| Mouse anti CEP164 | Santa Cruz Biotechnology | Cat # sc-515403, clone E9 |
| Mouse Anti-Arl13b | UC Davis/NIH NeuroMab Facility | Cat# 75-287, clone N295B/66 |
| Rabbit anti GFP | Invitrogen | Cat# A11122 |
| Rabbit anti ZO-1 | Invitrogen | Cat# 617300 |
| Rabbit anti Cone arrestin | Sigma-Aldrich | Cat# AB15282 |
| Mouse anti rhodopsin (RET-P1) | Santa Cruz Biotechnology | Cat# sc-57433, monoclonal |
| Mouse anti Vimentin | BD Transduction Labs | Cat# 550513, clone RV202 |
| Rabbit anti PKC-a | Sigma-Aldrich | Cat# P4334 |
| Rabbit anti TMEM119 | Abcam | Cat# ab209064 |
| Rat anti Adgre1 (F4/80) | Genetex | Cat# GTX26640, clone CI:A3-1 |
| Rabbit anti AIF1(IBA-1) | FUJIFILM (wako chemicals) | Cat# 019-19741 |
| Mouse anti Beta Catenin | BD Transduction Labs | Cat# 610154, clone 14/Beta-Catenin |
| Mouse anti aSMA | Millipore Sigma | Cat# A2547, clone 1A4 |
| IRDye® 800CW Goat anti-Rabbit IgG Secondary Antibody | Li-Cor Biosciences | Cat# 926-32211 |
| Goat anti-Rabbit IgG (H+L) Highly Cross-Adsorbed Secondary Antibody, Alexa Fluor™ Plus 488 | Thermo Fisher Scientific | Catalog # A-11034 |
| Goat anti-Mouse IgG (H+L) Highly Cross-Adsorbed Secondary Antibody, Alexa Fluor™ 594 | Thermo Fisher Scientific | Catalog # A-11032 |
| Donkey anti-Goat IgG (H+L) Highly Cross-Adsorbed Secondary Antibody, Alexa Fluor™ Plus 594 | Thermo Fisher Scientific | Catalog # A32758 |

| | | |
|---|---|---|
| Goat anti-Goat IgG (H+L) Highly Cross-Adsorbed Secondary Antibody, Alexa Fluor™ Plus 488 | Thermo Fisher Scientific | Catalog # A32723 |
| Alexa Fluor® 594 phalloidin | Thermo Fisher Scientific | Catalog # A12381 |
| Goat anti-Rat IgG (H+L) Highly Cross-Adsorbed Secondary Antibody, Alexa Fluor™ Plus 594 | Thermo Fisher Scientific | Catalog # A48264 |
| Goat anti-Mouse IgG (H+L) Highly Cross-Adsorbed Secondary Antibody, Alexa Fluor™ Plus 647 | Thermo Fisher Scientific | Catalog # A32728 |
| DAPI dihydrochloride | Sigma-Aldrich | D9542-1MG |

### Supplemental Figure 1

A

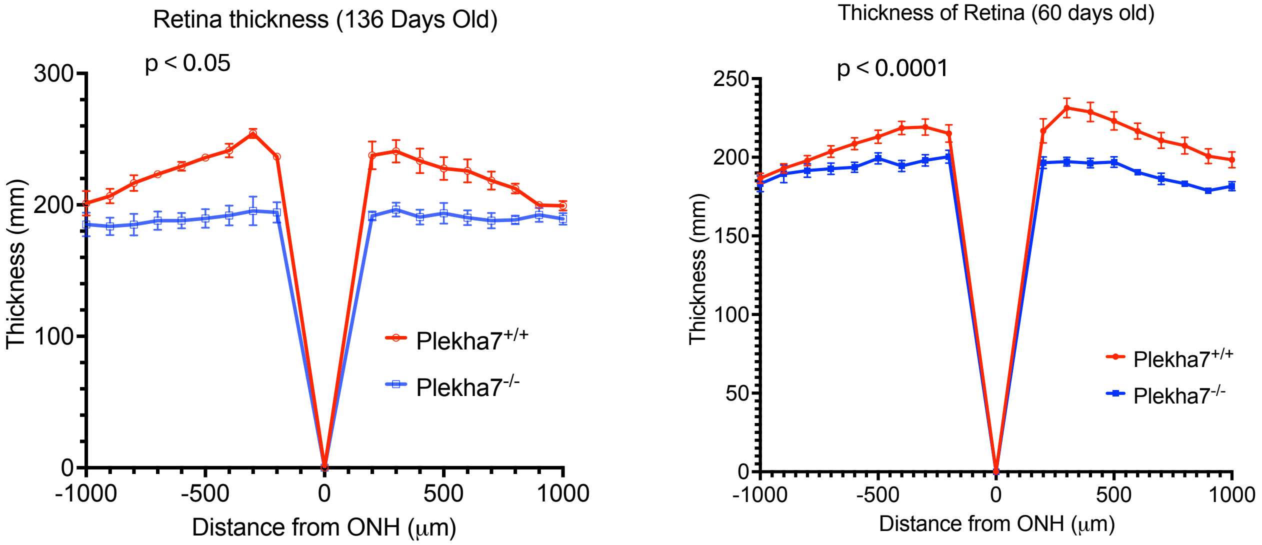


B

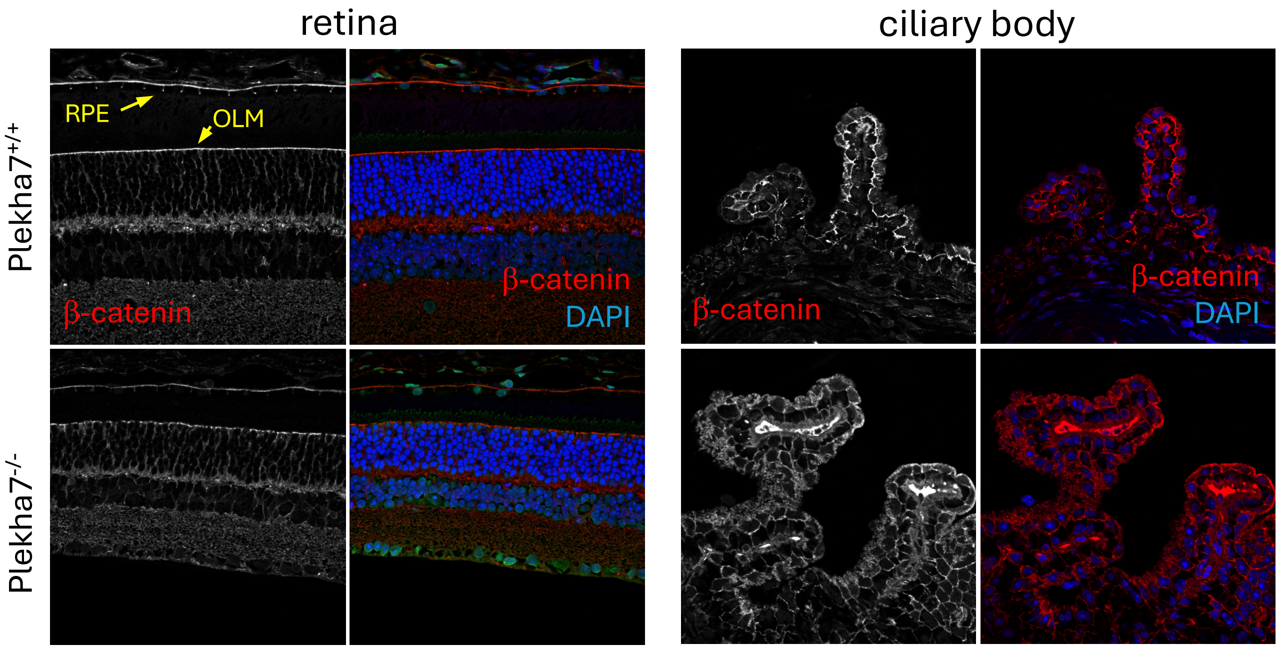

### Supplemental Figure 2

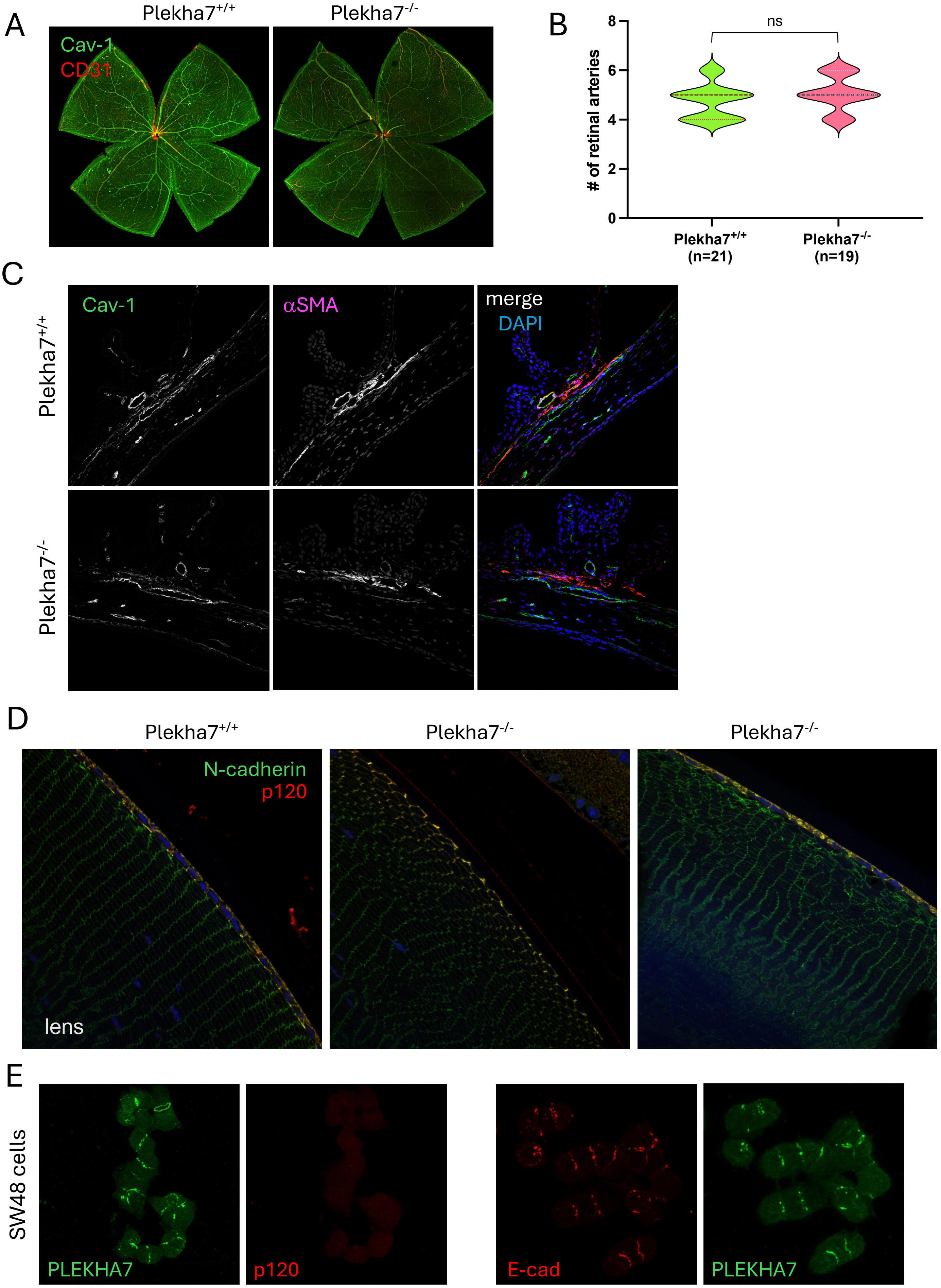

### Supplemental Figure 3

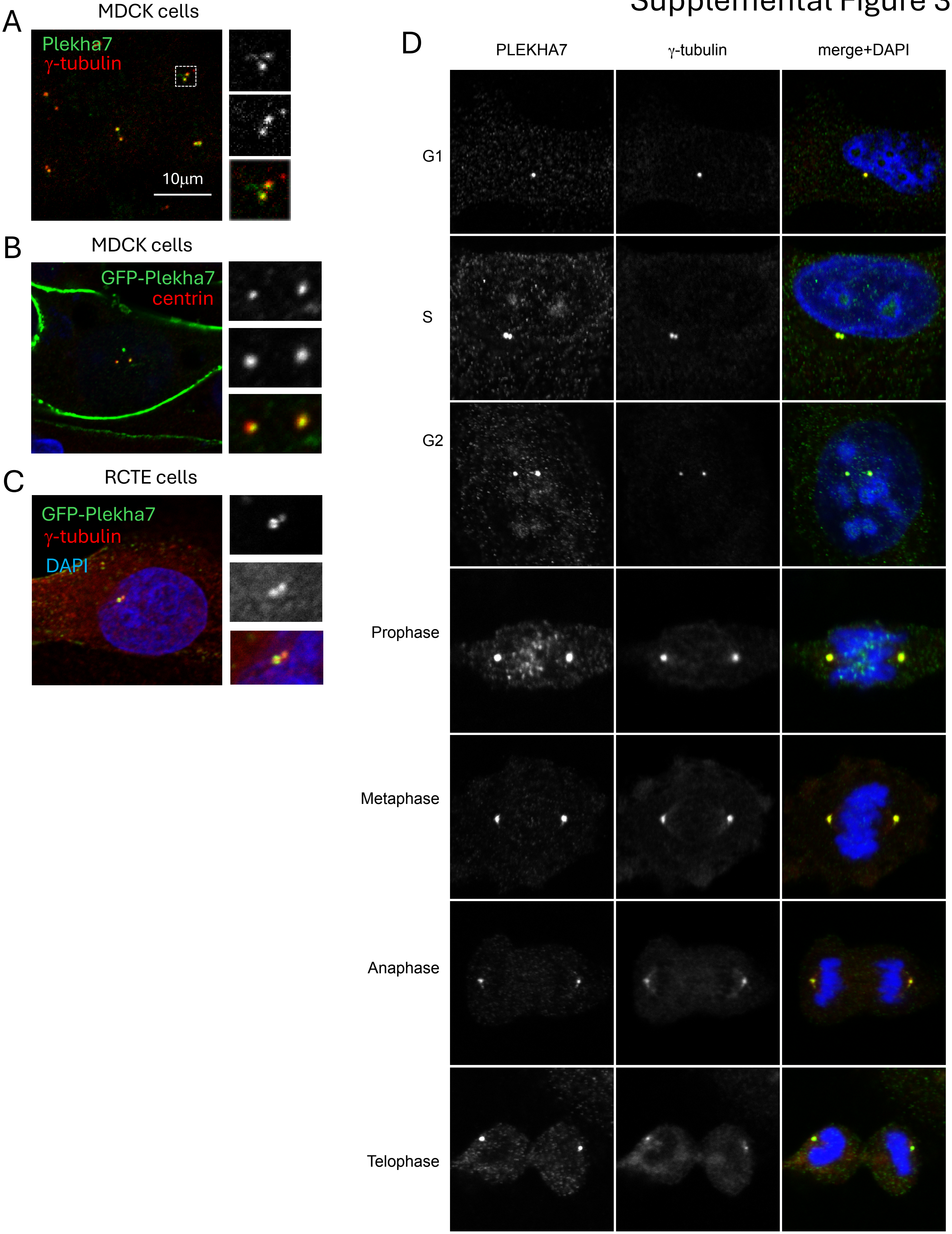

### Supplemental Figure 4

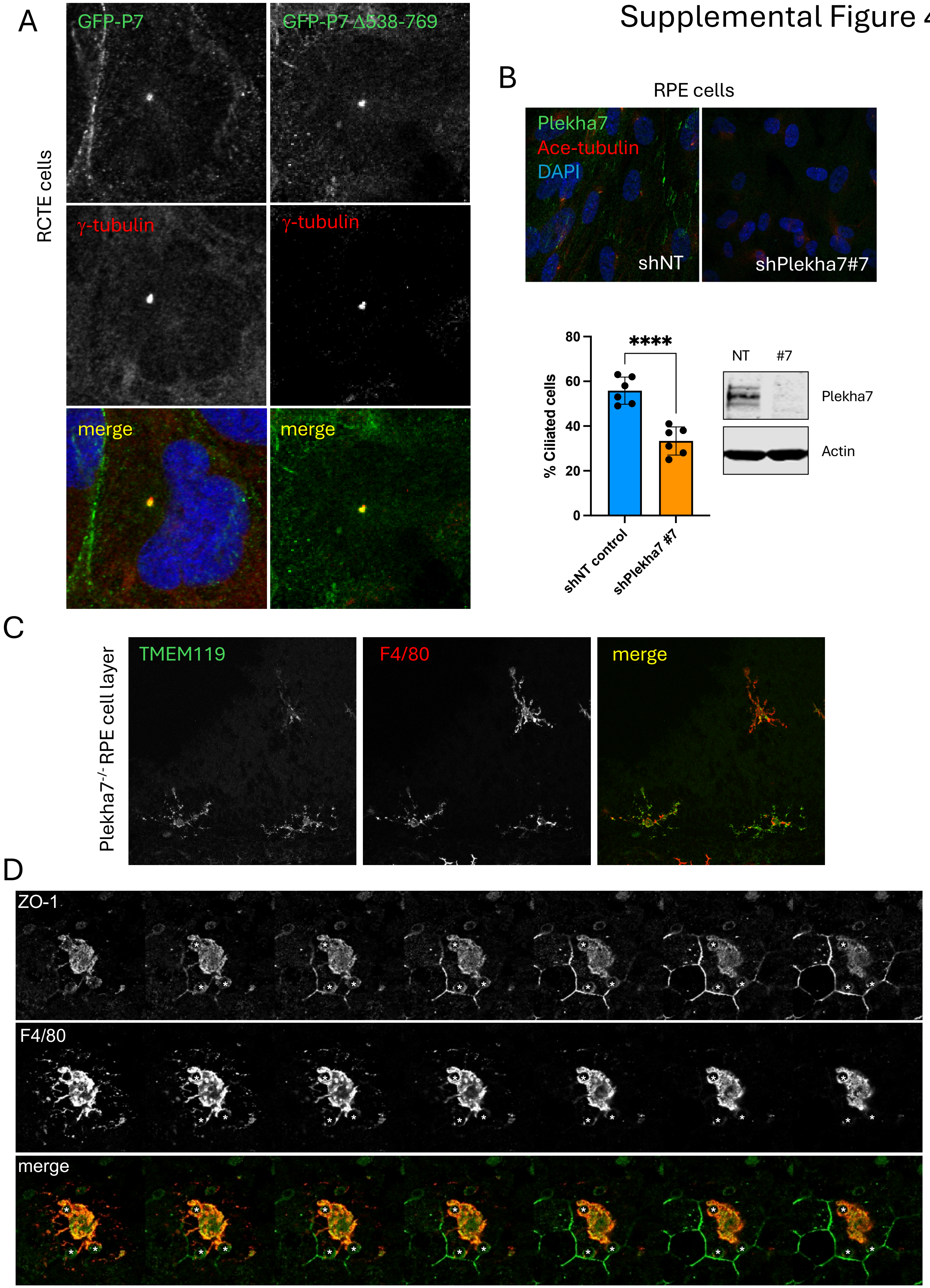
